# The *Salmonella* phage shock protein system is required for defense against host antimicrobial peptides

**DOI:** 10.1101/2025.04.16.649068

**Authors:** Marie-Ange Massicotte, Aline A. Fiebig, Andrei Bogza, Brian K. Coombes

## Abstract

Macrophages are professional phagocytes that play a major role in engulfing and eliminating invading pathogens. Some intracellular pathogens, such as *Salmonella enterica* serovar Typhimurium, exploit macrophages as niches for their replication, which requires precise and dynamic modulation of bacterial gene expression in order to resist the hostile intracellular environment. Here, we present a comprehensive analysis of the global transcriptome of *S.* Typhimurium across four stages of infection of primary macrophages. Our results revealed a profound change in early-stage gene expression dominated by pathways linked to metabolic processes required for *Salmonella* adaptation to the proinflammatory conditions of the macrophage. We identified the phage shock protein (Psp) system to be highly expressed in intracellular *S.* Typhimurium, with sustained high expression over the course of infection. We determined that the Psp system is regulated by the virulence-associated two-component system SsrA-SsrB, which coordinates its expression with critical bacterial functions required for immune evasion and intracellular survival. Functional assays demonstrated that the Psp system mediates resistance to host antimicrobial peptides, including cathelicidin-related antimicrobial peptide (CRAMP), which we demonstrate supports bacterial persistence in host tissues and survival within macrophages. Our findings establish the Psp system as a new and critical adaptive mechanism for evading host immune defenses and highlight the utility of temporal transcriptomics in unraveling the genetic strategies employed by *S.* Typhimurium during macrophage infection.

**Author summary:** *Salmonella enterica* is an important global pathogen that infects a wide range of mammalian hosts, requiring it to survive in diverse and hostile environments. A key aspect of *Salmonella* pathogenesis is its ability to reside within host immune cells like macrophages, where it must rapidly adapt to intracellular conditions. To do so, the bacteria must ensure correct spatiotemporal expression of virulence genes to maximise fitness in each environment. To better understand how *Salmonella* modulates its gene expression during infection of host cells, we defined its transcriptome at four distinct stages of primary murine macrophage infection. Our findings reveal that the first stage of early infection is dominated by changes in gene expression of metabolic circuits. Furthermore, we identified the phage-shock protein (Psp) system as highly expressed within intracellular *Salmonella* throughout the course of infection as a result of regulatory evolution that coordinates its expression with virulence genes. We showed that this system is required for bacterial survival within macrophages and host tissues by mediating resistance to host cationic antimicrobial peptides. These findings highlight the dynamic nature of *Salmonella’s* transcriptional response during macrophage infection and uncover a previously unknown function of the Psp system as an adaptive mechanism for evading host immune defenses.

## Introduction

*Salmonella enterica* serovar Typhimurium (*S.* Typhimurium) is a leading cause of gastrointestinal disease worldwide [1]. Although enteric salmonellosis is generally self-limiting, infections can escalate to life-threatening systemic illnesses in immunocompromised individuals or when caused by emerging invasive non-typhoidal *Salmonella* (iNTS) strains [2,3]. The efficacy of antibiotic therapy has been consistently compromised by the emergence of multidrug resistant strains [4,5]. Addressing this global health challenge requires a deeper understanding of the virulence mechanisms underlying *S.* Typhimurium pathogenesis to identify novel targets for anti-*Salmonella* therapies.

A hallmark of *S.* Typhimurium pathogenesis is its capacity to evade the innate immune response and establish a replication niche within innate immune cells, particularly macrophages. Intracellular replication occurs within a specialized phagosome called the *Salmonella*-containing vacuole (SCV) [6] where *Salmonella* resists phagosomal destruction while accessing nutrients to support replication and survival [7,8]. Infected macrophages can serve as reservoirs for bacterial dissemination to systemic sites [5]. Despite offering a replicative niche, the SCV presents several antimicrobial challenges, including an acidified microenvironment, restriction of essential ions and metals, and exposure to toxic compounds such as cationic antimicrobial peptides (cAMPs), reactive oxygen species (ROS), and reactive nitrogen species (RNS) [9]. To overcome these barriers, *S.* Typhimurium employs tightly regulated gene networks that integrate signals originating from the innate immune response with transcriptomic changes necessary for intracellular survival [10–12].

Despite decades of study, the mechanisms underlying *S.* Typhimurium’s evasion of host defenses and the gene networks that underpin these processes remain incompletely understood. Early microarray studies provided foundational insights into *S.* Typhimurium gene expression during macrophage or epithelial cell infections [13,14]. However, these approaches lacked the resolution that deep-sequencing technologies now offer [15]. RNA sequencing (RNA-seq) has been used to define the primary transcriptome of *S.* Typhimurium during macrophage infection, but these studies have relied on single time-point analyses [16], precluding an understanding of the temporal transcriptional response. Furthermore, most intracellular transcriptomics studies have used immortalized cell lines such as RAW264.7, J774A.1, or HeLa cells, which only partially recapitulate the biological processes occurring in primary cells [17]. Investigations using primary cell models are required to uncover the transcriptional strategies employed by *S.* Typhimurium in biologically relevant contexts [18].

Here, we examined the transcriptional response of *S.* Typhimurium at four distinct stages of infection in primary bone marrow-derived macrophages. We found that the most extensive transcriptional changes occurred during the early stages of infection, particularly within the first four hours. Among the genes most highly upregulated during this period, the phage shock protein (Psp) system, an envelope stress response pathway critical for mitigating inner membrane damage, emerged as a key transcriptional response. We demonstrated that the major phenotypic output of Psp system activation is bacterial resistance to host antimicrobial peptides and survival within macrophages, and that this regulation appears to have been selected via cis-regulatory evolution. Together, these results provide new insights into the temporal transcriptional adaptations of *S.* Typhimurium during macrophage infection and establish a novel role for the Psp system in bacterial resistance to host antimicrobial defenses.

## Results

### Intracellular *S.* Typhimurium undergoes rapid transcriptional changes during the early stages of macrophage infection

The macrophage niche is a highly dynamic environment to which *S.* Typhimurium must continuously adapt for intracellular survival. To assess the extent of transcriptional changes in intracellular *S.* Typhimurium, we analyzed its transcriptome at four stages of infection in primary macrophages that we defined as onset, early, middle, and late (Fig 1A). The stages were based on previous literature [13] and reflect key events encountered by *S.* Typhimurium within the SCV, including phagosome acidification [19], oxidative and nitrosative stress [20,21], SCV maturation [22], and exposure to cAMPs [23]. Bone marrow-derived macrophages (BMDMs) from C57BL/6J mice were infected with wild-type *S.* Typhimurium, and RNA from intracellular bacteria was extracted at 0, 4, 8, and 12 h post-infection [13,16]. cDNA libraries reached a sequencing depth of ∼ 20 million reads that mapped to the *Salmonella* reference genome for ∼ 600x genome coverage for each library [24,25].

**Fig 1.**
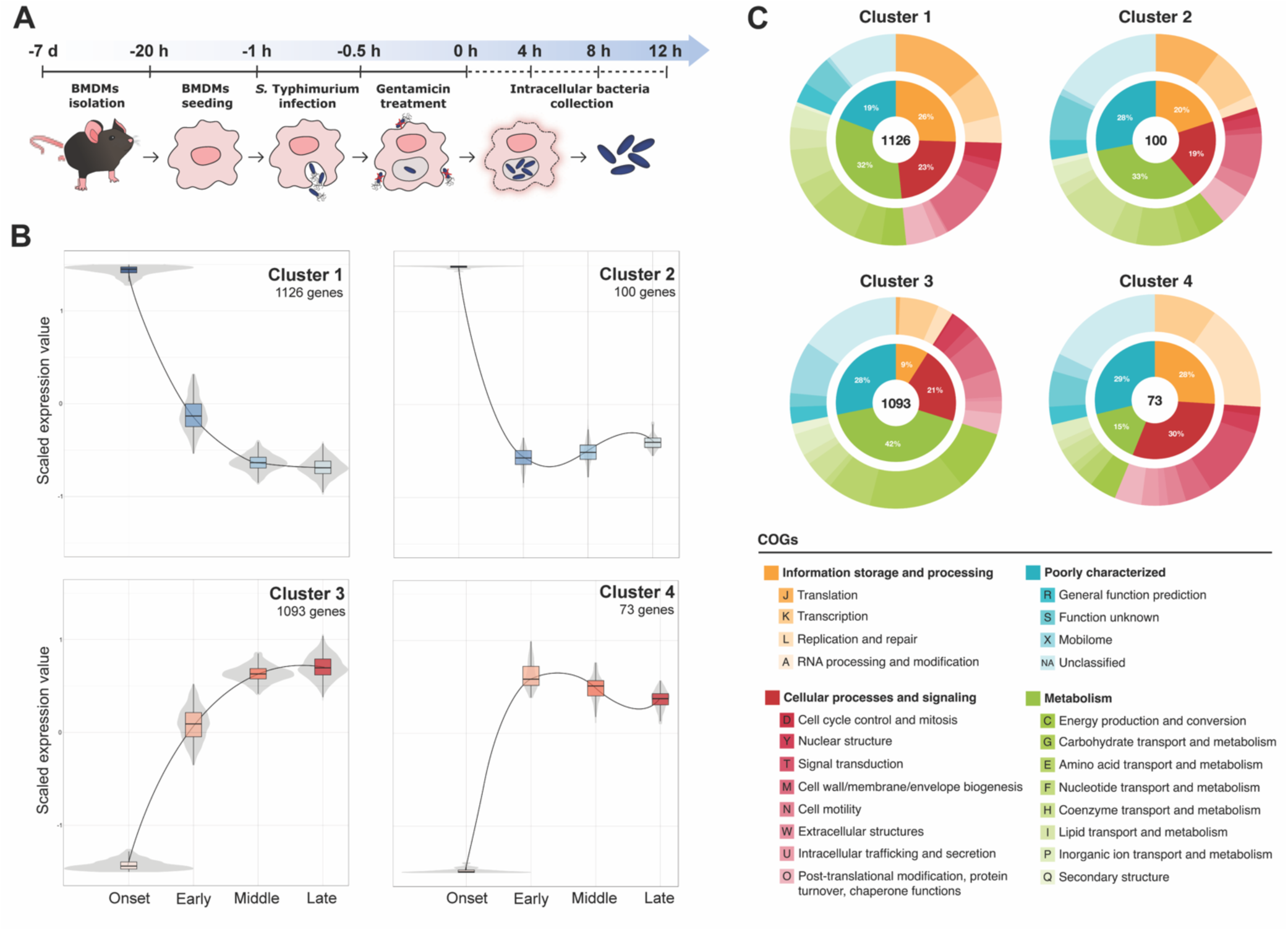
*S.* Typhimurium transcriptional profile changes drastically between onset and early stages of macrophage infection. (A) Schematic representation of the experiment. (B) Clusters of significantly differentially regulated genes in S. Typhimurium during the infection of macrophages identifies four patterns of expression over time. Genes are plotted on the y-axis according to a scaled expression value (z-score). For each cluster, volcano and box plots were constructed for each time point, with median expression level denoted by the horizontal line. Lines connect the mean expression levels of consecutive time points, to display the gene expression trend of that cluster. Genes that did not match the expression profiles of any of the clusters were omitted. Clustering was performed using DEGreport with genes of an adjusted P value < 0.01 calculated by the LTR test of DESeq2. (C) All differentially expressed genes identified in (B) were assigned to a COG functional category. Percentages reflect the abundance of each major COG category (inner circle) in each cluster.

To identify *S.* Typhimurium genes with significant changes in expression across the different stages of infection, we performed differential expression analysis using the Likelihood ratio test (LRT) in DESeq2. This method, ideal for comparative analysis within a time series [26], identified 2,386 differentially expressed genes throughout macrophage infection, representing nearly half of the *S.* Typhimurium genome. Clustering analysis of significant genes revealed four expression patterns, the majority of which fell into Cluster 1, representing peak expression at onset (1,126 genes) or Cluster 3 representing peak expression at late infection (1,093 genes) (Fig 1B, S1 Table). These data highlight a striking transcriptional shift during the transition from onset to early infection, suggesting a crucial period of widespread genetic reprogramming as an adaptive response to the intracellular environment. Building on previous microarray studies [13], these data indicate that early infection represents a pivotal stage of S. Typhimurium adaptation and survival within host macrophages.

To investigate the biological processes reflected within each expression cluster, genes were assigned to clusters of orthologous groups (COGs) categories (Fig 1C, S1 Table). Genes with decreasing expression over time (clusters 1 and 2) were predominantly linked to energy-intensive processes and rapid growth. For example, we observed decreased expression of genes involved in ribosome biogenesis, including the *rps* and *rpl* operons [27], the *rsgA* and *rim* operons encoding accessory proteins involved in ribosome assembly and maturation, and the *rsm* genes responsible for rRNA modifications. In line with this, genes involved in cell division, *fts* and *zip*, shared this expression pattern [28]. Interestingly, several ROS detoxifying enzymes, including *katG*, *sodA*, *sodCII*, and *ahpC* [29], grouped into expression clusters 1 and 2, suggesting that *S.* Typhimurium resists an early respiratory burst that subsides as the infection progresses. Together, these data indicated that *S.* Typhimurium transitions from early, active replication to energy conservation that supports survival in the host macrophage.

In contrast with this metabolic reprogramming, our data revealed that genes whose expression increases over time (clusters 3 and 4) are predominantly associated with alternative respiratory metabolism and adaptation to nutrient scarcity. Genes with dominant expression in these categories included enzymes required to sustain anaerobic respiration, such as the nitrate reductases *nar* and *nap* operons [30], the *ttr* operon encoding a tetrathionate reductase [31], the *hyp* operon encoding for an electron donor NiFe hydrogenase for fumarate respiration mediated by a fumarate reductase encoded by *frdABCD* [32], and the *dmsABC* operon which encodes for the three subunits of the anaerobic dimethyl sulfoxide reductase [33]. Additionally, we detected the upregulation of several genes involved in lipid and sugar metabolism previously shown to be important for replication in mice and macrophages [34,35]. This included the *eut* operon, required for the utilization of ethanolamine as a source of carbon and nitrogen and as an environmental cue used by *S.* Typhimurium to coordinate metabolism and virulence [36,37]. We also detected an upregulation of the *pdu* operon required for the degradation of 1,2-propanediol, the end-product of the anaerobic fermentation of fucose and rhamnose [38]. The genes involved in the degradation of fucose (*fuc* operon) and rhamnose (*rha* operon) were also upregulated in the late infection clusters [39]. Genes encoding components of the phosphotransferase system involved in the import and phosphorylation of various sugars like glucose, galactitol, mannitol, and fructose were upregulated [34]. These data suggest that in response to the oxygen and nutrient-depleted SCV, *S.* Typhimurium shifts toward anaerobic respiration via different terminal electron acceptors to support growth using host-derived nutrients as energy sources. Taken together, our temporal transcriptomic analysis highlights the importance of metabolic plasticity for *S.* Typhimurium to adapt and survive within the intracellular environment during macrophage infection.

### Time-course RNA sequencing reveals upregulation of the *psp* operon by intracellular *S.* Typhimurium

Our clustering analysis revealed that *S.* Typhimurium undergoes substantial transcriptional changes during the onset and early stages of macrophage infection. To quantify these changes, we performed differential expression analysis of *S.* Typhimurium genes at early, middle, and late stages of infection relative to the onset stage using DESeq2 (Fig 2A and S2 Table) [26]. Approximately 50% of *S.* Typhimurium genes were differentially regulated in at least one stage of infection compared to onset, consistent with the results of the clustering analysis. Screening operons for genes that were significantly upregulated across all stages of infection following infection onset identified the entire phage shock protein (*psp*) system as highly expressed throughout infection, suggesting a pivotal role in supporting intracellular infection (Fig 2B).

**Fig 2.**
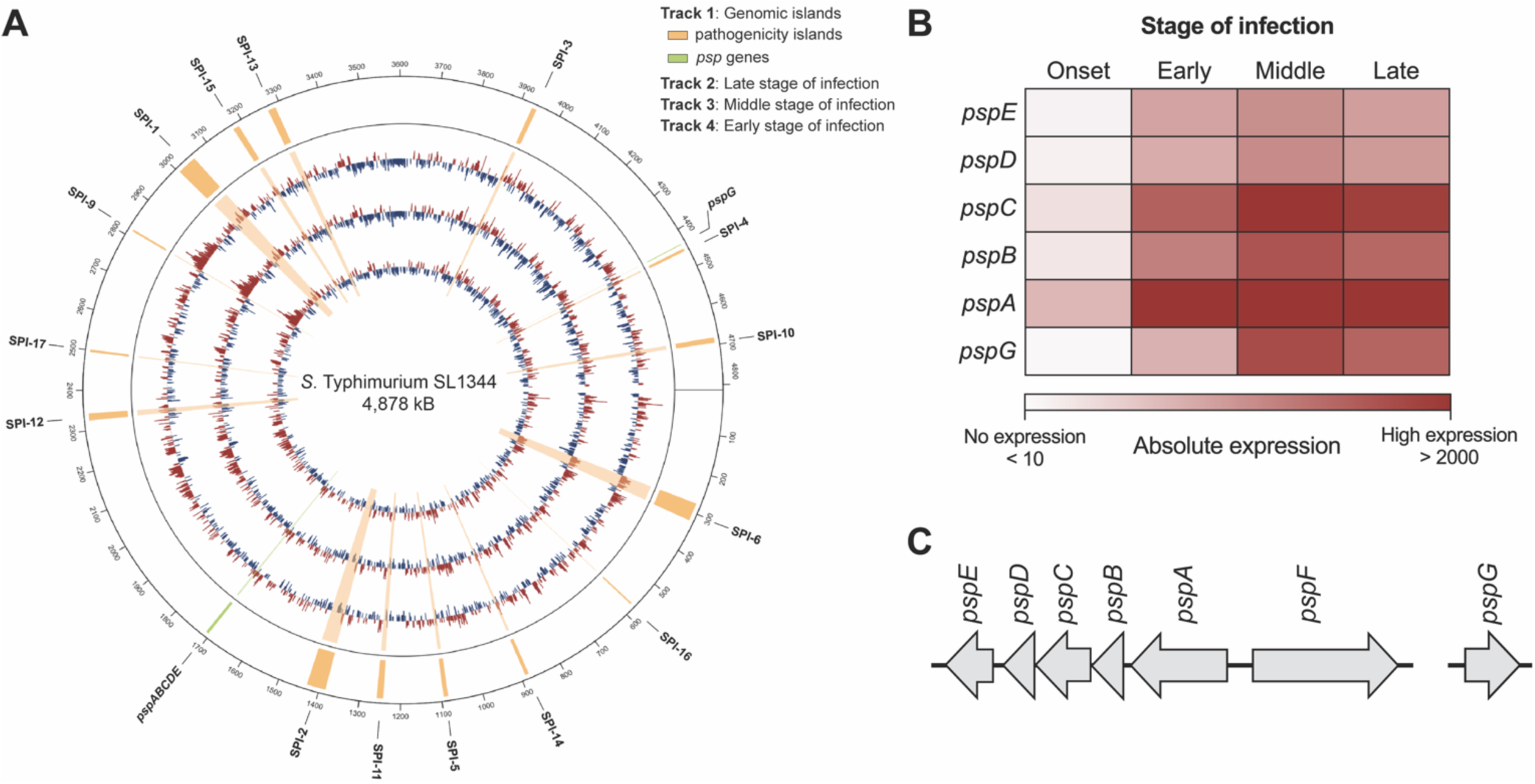
The *S.* Typhimurium *psp* operon is upregulated during the infection of macrophages. (A) Circos plot representing the genome-wide fold change in *S.* Typhimurium gene expression at early, middle, and late stages of infection relative to the onset stage from the inside to the outside tracks respectively. Red peaks indicate significantly up-regulated genes (log2 fold change >2), blue peaks indicate significantly down-regulated genes (log2 fold change >-2). Genes deemed not significantly regulated are not depicted. Genomic islands are indicated and labelled on track 1 (outermost circle), where pathogenicity islands are in orange and *psp* genes are in green. (B) Heatmap showing expression of S. Typhimurium *psp* genes that are upregulated during the onset, early, middle, and late stages of macrophage infection. The heatmap colors represent the absolute expression levels (TPM values) based on the color bar below. (C) Schematic representation of the *psp* operon in S. Typhimurium.

The Psp system is an envelope stress mechanism encoded by the *psp* operon (*pspABCDE*), the unlinked *pspG* gene, and the transcriptional activator *pspF* located upstream of *pspA* (Fig 2C). Studies in *Escherichia coli* have shown that the Psp system is activated by damage to the inner membrane or loss of the proton motive force (PMF). Upon induction, PspA preserves the PMF, while PspB and PspC mitigate toxicity from mislocalized secretins [40,41]. Currently, little is known about the biological functions of the other Psp proteins. Although the expression of *psp* genes in *S.* Typhimurium has been reported during infection of epithelial cells and macrophages [13,14,16], the physiological significance of this upregulation for its pathogenesis has not been elucidated.

### The transcriptional regulator SsrB controls the expression of the Psp system

*S.* Typhimurium relies on a complex regulatory network of two-component regulatory systems (TCSs) to coordinate the expression of virulence genes essential for intracellular survival [42]. Given the strong upregulation of *psp* genes during macrophage infection, we considered the possibility that the Psp system in *Salmonella* was regulated by a TCS active in the intracellular niche. Specifically, we investigated PhoQ-PhoP, PmrB-PmrA, and SsrA-SsrB, which are known to regulate *S.* Typhimurium’s intracellular fitness and virulence features [43–45]. To assess the involvement of these TCS, we constructed a luciferase transcriptional reporter of the *pspABCDE* promoter (P*pspABCDE*-lux) and monitored promoter activity over 16 h in wild-type *S.* Typhimurium and strains with deletions of *phoP*, *pmrA*, or *ssrB* (Δ*phoP*, Δ*pmrA*, Δ*ssrB*). Deletion of *phoP* or *pmrA* significantly increased *psp* promoter activity compared to wild type (Fig 3A), consistent with earlier findings in a *rpoE* mutant [46]. In contrast, the deletion of *ssrB* completely abolished *psp* promoter activity, establishing the virulence-associated SsrA-SsrB TCS as a critical regulator of the Psp system.

**Fig 3.**
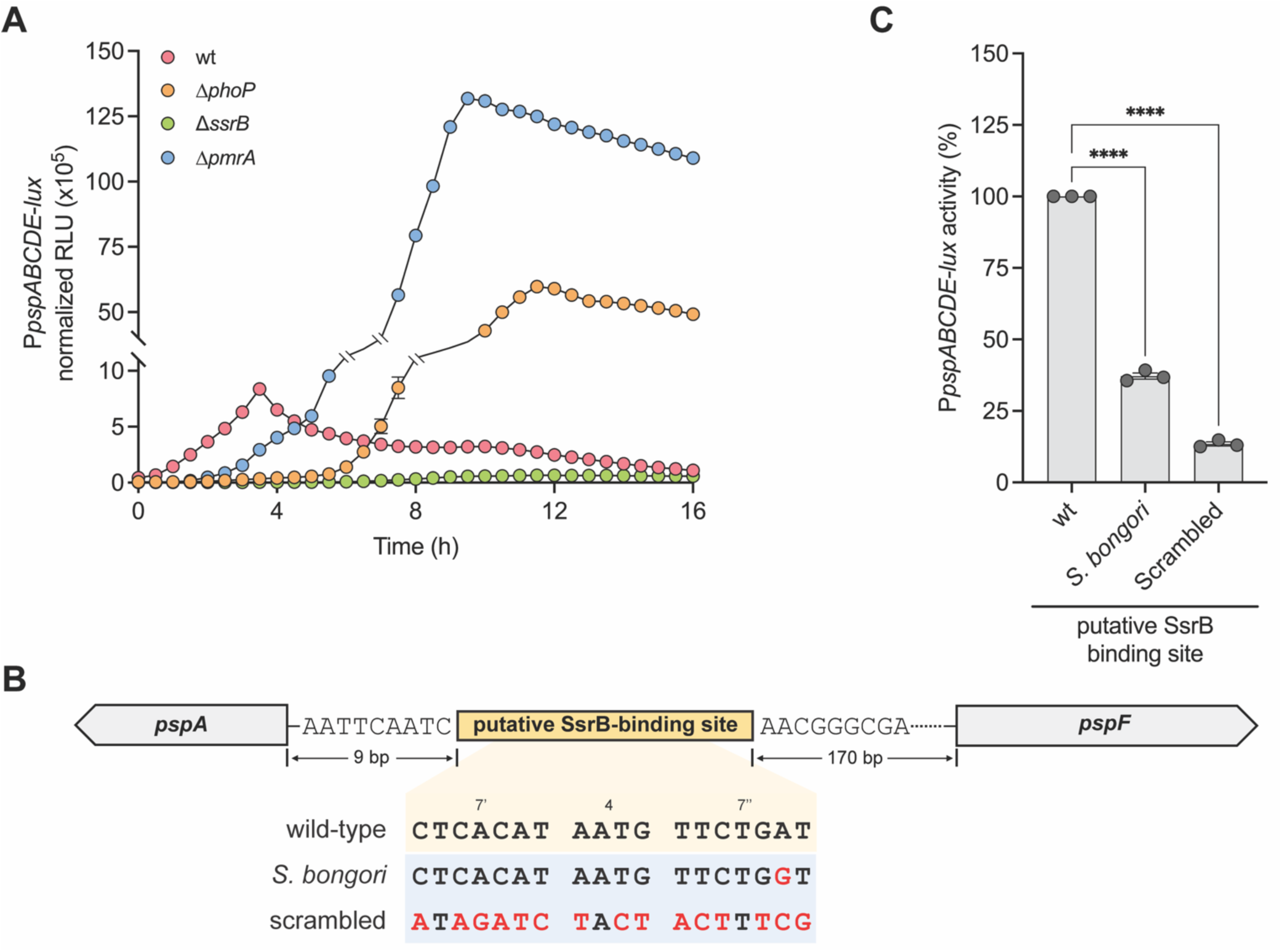
The Psp system is transcriptionally regulated by SsrB. (A) Transcriptional reporter of the *pspABCDE* operon promoter in Δ*phoP*, Δ*pmrA*, Δ*ssrB*, or wild-type (wt) *S.* Typhimurium strains. Data are mean relative light units (RLU) normalized to the optical density of the culture from three independent experiments. (B) Graphical representation of the putative SsrB-binding site identified upstream of the *pspA* gene. The sequences of the wild-type SsrB-binding motif and mutated binding sites from the *psp* promoter are shown. The bases colored in red are used to highlight the mutated bases in putative *S. bongori* and scrambled SsrB-binding site relative to wild-type. (C) Transcriptional reporter data for the wild-type (wt) and the two mutated SsrB-binding site as defined in (B). Promoter activity was measured as the ratio of RLU/optical density normalized to the promoter activity from the wild-type reporter at 4 h. Data are the means standard error of the mean from three independent experiments. Groups were compared against wild-type via one-way ANOVA. ****p < 0.0001 (Holm-Sidak’s multiple comparisons test).

SsrB has been shown to capture a large genetic network in *S.* Typhimurium that integrates both ancestral and horizontally acquired genes via regulatory evolution [47,48]. To determine if the Psp system in *Salmonella* has undergone regulatory evolution and selection for SsrB control, we scanned the intergenic region upstream of *pspA* for evidence of the flexible 18 bp palindrome sequence that defines DNA recognition by SsrB (S3 Fig). This analysis identified a putative SsrB binding site 9 bp upstream of the transcriptional start site of *pspA* (Fig 3B). To validate this site as an input sequence for SsrB regulation, we designed two transcriptional reporters; one replacing the SsrB binding site with the homologous sequence from *S. bongori,* which lacks the SsrA-SsrB TCS but contains an otherwise identical *psp* operon, and a second reporter with a scrambled version of the putative SsrB binding sequence in which the 18 bp palindrome was randomized (Fig 3B). The strain containing the *S. bongori* replacement sequence, with only a single nucleotide replacement, reduced *psp* promoter activity to ∼35% of wild-type levels, whereas the scrambled binding site produced only ∼12% residual activity relative to wild-type (Fig 3C and S3 Fig). Taken together, these data established that the Psp system is regulated by the SsrA-SsrB TCS in *S.* Typhimurium and implied an important role for this system in intracellular survival in macrophages.

### The Psp system promotes *S.* Typhimurium survival in primary macrophages and contributes to its persistence in host tissues

The upregulation of *psp* genes during macrophage infection, and the apparent regulatory evolution of this system for SsrB control, strongly suggested that the Psp system plays a critical role in intracellular *S.* Typhimurium pathogenesis. To assess whether Psp deficiency affected bacterial survival in macrophages, we infected BMDMs isolated from C567BL/6J mice with either wild-type S. Typhimurium or an isogenic mutant with a deletion of *pspA* (Δ*pspA*) and monitored bacterial survival over time. In the first 12 h after infection, there was a 200% increase in intracellular wild-type bacteria, whereas survival of the Δ*pspA* mutant was only ∼50% and significantly decreased further to 30% after 24 h of infection (Fig 4A). These findings indicate that PspA is required for *S.* Typhimurium survival and replication in primary macrophages, consistent with the elevated expression of the Psp system during intracellular infection.

**Fig 4.**
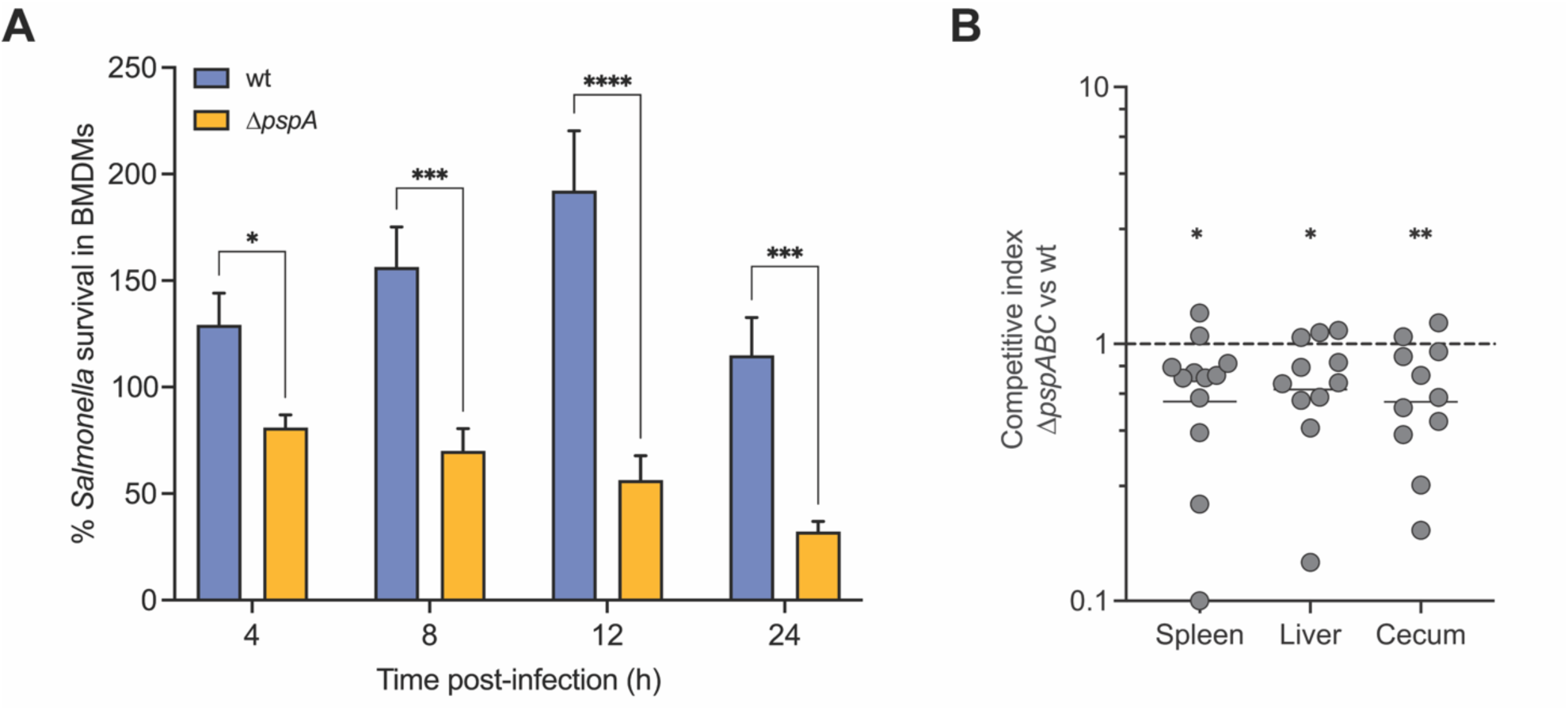
Psp-deficient *S.* Typhimurium bacteria are attenuated for virulence in primary macrophages and *in vivo*. (A) *S.* Typhimurium Δ*pspA* bacteria show increased susceptibility to killing by primary macrophages compared to wild-type (wt) bacteria. Survival was measured as the intracellular bacterial burdens enumerated at 4, 8, 12, and 24 h following infection normalized to the values of the initial number of internalized bacteria (T0). Bar plots depict mean of at least four biological replicates and error bars indicate standard error of the mean. Groups were compared by two-way ANOVA, *p < 0.05, ***p < 0.001, ****p < 0.0001 (Holm-Sidak’s multiple comparisons test). (B) Wild-type *S.* Typhimurium out-competed the Δ*pspA* mutant in all gut tissues. C57BL/6J mice were infected by intraperitoneal injection with equal number of wild-type *S.* Typhimurium and Δ*pspA* mutant strains, and the competitive index was calculated after 2 days of infection. Each data point represents the value for an individual animal, and the horizontal lines indicate geometric means. The broken line shows a competitive index of 1, representing equal fitness. Groups were compared against a value of 1 using one-sample parametric T-test, *p < 0.05, **p < 0.01. Data are from two independent experiments.

To evaluate whether the Psp system contributes to *S.* Typhimurium fitness during *in vivo* infection, we constructed a Δ*pspABC* mutant strain in SL1344 and assessed its infection potential in C57BL/6J mice. To do this, we performed competitive infections between the Δ*pspABC* mutant and wild-type bacteria and quantified bacterial burden two days after infection in the spleen, liver, and cecum. In all three tissues, wild-type bacteria significantly out-competed the Δ*pspABC* mutant (CI, spleen; 0.7043, P < 0.05, CI liver; 0.7453, P < 0.05, and CI cecum; 0.6779, P < 0.01) (Fig 4B). These results are consistent with those of a previous report [49], and demonstrate that Psp-deficient *S.* Typhimurium is attenuated for virulence *in vivo*. Together, these results established that the Psp system contributes to bacterial fitness in both primary macrophages and mouse models of infection.

### PspA mediates resistance to the murine antimicrobial peptide CRAMP

To investigate how the Psp system supports *S.* Typhimurium survival during infection, we focused on the functional role of PspA. In *E. coli*, the Psp system, particularly PspA, is thought to preserve the PMF during extracytoplasmic stress [50,51]. In *S.* Typhimurium, while the physiological role of Psp proteins remains poorly understood, PspA has been implicated in divalent metal transport [52], resistance to the proton ionophore CCCP, and survival under stress conditions such as stationary phase and resistance to the innate immune protein bactericidal permeability-increasing protein (BPI) in *S.* Typhimurium lacking the alternative sigma factor RpoE [46]. Given the association of these conditions with PMF disruption, we first tested the susceptibility of a Δ*pspA* mutant to PMF-disrupting agents [53,54]. For this, we determined the minimum inhibitory concentrations (MIC) of polymyxin B, colistin (polymyxin E), and CCCP for the wild-type and Δ*pspA* mutant strains in acidic, low-phosphate, low-magnesium media (LPM) that was established to mimic conditions of the SCV [16,55,56]. The Δ*pspA* mutant showed significantly increased susceptibility compared to the wild-type, with a 16-fold, 32-fold, and 4-fold increase in susceptibility to polymyxin B, colistin, and CCCP, respectively (Fig 5A). To confirm that the mutant’s susceptibility was specific to PMF disruption and not general membrane damage, we tested MICs of EDTA and polymyxin B nonapeptide (PMBN), which both disrupt the outer membrane without causing direct damage to the inner membrane [57,58]. No differences in MICs were observed between the wild-type and Δ*pspA* mutant for these agents (Fig 5A), confirming that PspA confers resistance to PMF-disrupting agents likely through inner membrane destabilization.

**Fig 5.**
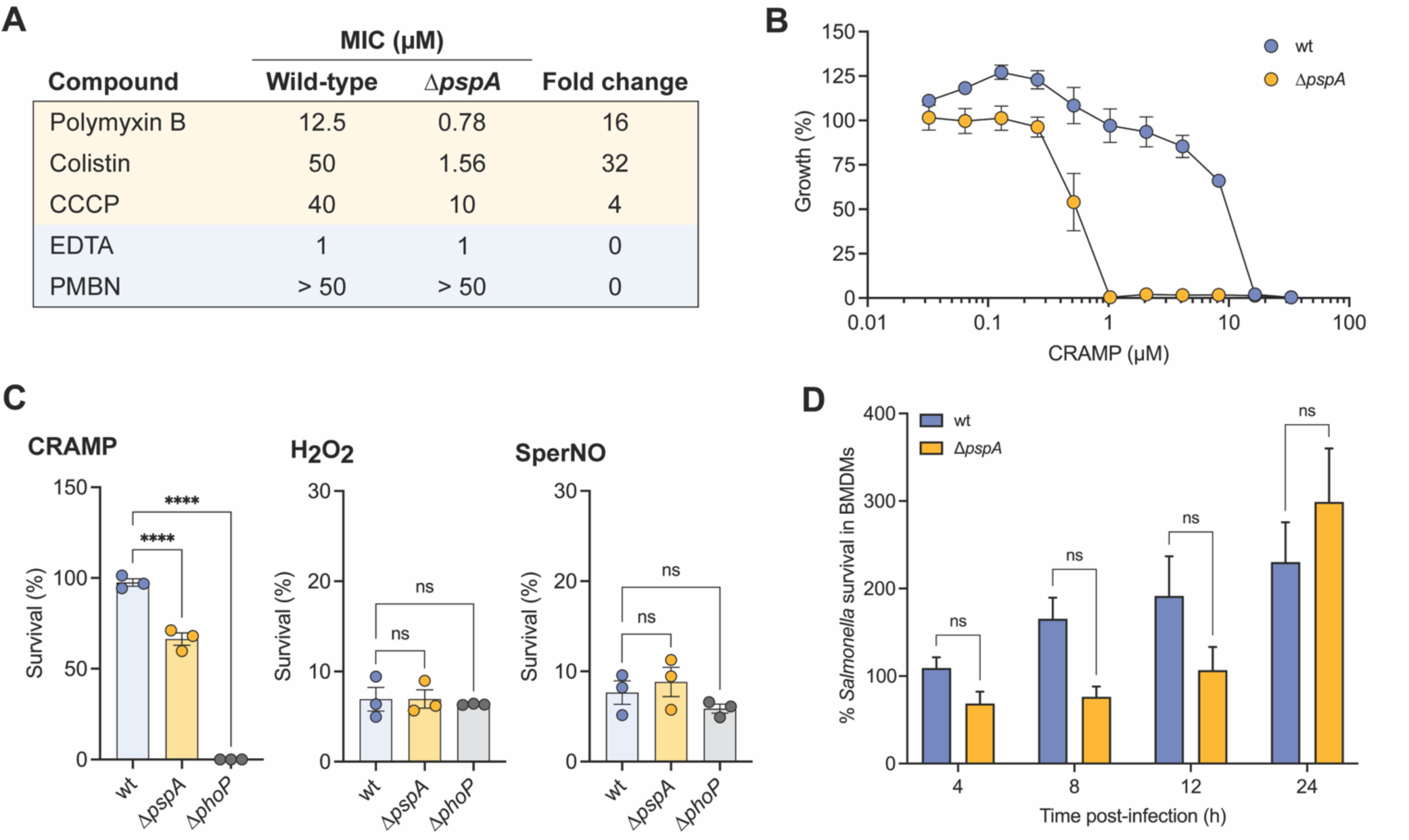
PspA is required for resistance to CRAMP. (A) Table of MIC values for various PMF disruptors (polymyxin B, colistin, and CCCP) and outer-membrane disruptors (EDTA, and polymyxin B nonapeptide; PMBN) against wild-type *S.* Typhimurium and Δ*pspA* mutant in infection-mimicking media. The Δ*pspA* mutant shows increased susceptibility to all the PMF disruptors tested relative to wild-type, but not the outer-membrane disruptors. Data are representative of three biological replicates. (B) Growth of wild-type *S.* Typhimurium and Δ*pspA* mutant in the presence of CRAMP in infection-mimicking media. Data is from three biological replicates, dots and error indicate mean and standard error of the mean. (C) *S.* Typhimurium Δ*pspA* mutant is more susceptible to killing by CRAMP than the wild-type (wt) is, but not to hydrogen peroxide (H_2_O_2_) nor Spermine NONOate (SperNO) in infection-mimicking media. Bacteria were incubated with 10μM of CRAMP or 50μM of H_2_O_2_, or 50μM of SperNO for 3 h. *S.* Typhimurium Δ*phoP* mutant was used as a control for extreme sensitivity to antimicrobial peptides. Bar plots depict mean of three biological replicates and error bars indicate standard error of the mean. Groups were compared against wild-type via one-way ANOVA. ****p < 0.0001 (Holm-Sidak’s multiple comparisons test); ns, not significantly different. (D) Intracellular replication defect of *S.* Typhimurium Δ*pspA* bacteria is abrogated in *Camp*^-/-^ macrophages. As in Fig 3A, survival was measured as the ratio of the number of viable bacteria enumerated at 4, 8, 12, and 24 h compared to the initial number of internalized bacteria at T0. Bar plots depict mean of three biological replicates and error bars indicate standard error of the mean.

Next, we investigated the protective role of PspA against host-derived antimicrobial factors. Among these, cationic antimicrobial peptides (cAMPs) are important host defense mechanisms that target the bacterial cytoplasmic membrane to induce cell lysis [59–62]. In murine macrophages, the primary cAMP is cathelicidin-related antimicrobial peptide (CRAMP), a homolog of human LL-37 [63]. Infections with different intracellular bacteria have been shown to induce CRAMP upregulation [64,65], and CRAMP colocalizes with the SCV during *S.* Typhimurium infection [23]. MIC assays revealed that the Δ*pspA* mutant was 16-fold more susceptible to CRAMP than wild-type (Fig 5B). Complementation of the Δ*pspA* mutant with *pspA* expressed from its native promoter completely restored resistance to CRAMP to wild-type levels (S2 Fig). Survival assays further demonstrated Δ*pspA* mutant susceptibility to CRAMP. After exposure to 10 μM CRAMP for 3 hours, ∼60% of the Δ*pspA* mutant bacteria survived compared to nearly 100% survival for wild-type (Fig 5C). As a control, we included a Δ*phoP S.* Typhimurium mutant which has extreme sensitivity to cAMPs [66,67]. As expected, we observed the complete killing of the Δ*phoP* strain after 3 hours of CRAMP exposure. These results establish that PspA-deficient *S.* Typhimurium has a significant survival defect in the presence of CRAMP. To probe if PspA is important for resistance against other key host-derived antimicrobial factors found within the SCV, we repeated the survival assays in the presence of oxidative and nitrosative stressors. Survival assays using hydrogen peroxide and nitric oxide showed no significant differences between the Δ*pspA* mutant and wild-type (Fig 5C), suggesting that PspA’s protective role is more specific to AMP resistance.

Finally, to validate that PspA promotes *S.* Typhimurium survival within host macrophages by promoting resistance to CRAMP, we assessed bacterial survival in macrophages lacking CRAMP harvested from *Camp*^-/-^ mice. In contrast to infections in CRAMP-producing macrophages from C57BL/6J mice (Fig 4A), Δ*pspA* mutants showed no survival defect in *Camp*^-/-^ macrophages, with both mutant and wild-type strains exhibiting over 200% increase in viable bacteria after 24 hours (Fig 5D). These findings confirm that the fitness defect of PspA-deficient *S.* Typhimurium in macrophages is specifically linked to increased susceptibility to CRAMP.

## Discussion

The ability to precisely regulate gene expression in response to environmental cues is essential for *S.* Typhimurium’s intracellular survival, enabling adaptation to hostile immune challenges within the macrophage niche. Despite the importance of these transcriptional changes, which have mostly been defined at single time points during infection, less is known about the dynamic transcriptional response of *S.* Typhimurium over the course of infection in primary macrophages. By employing RNA sequencing to profile the bacterial transcriptome at four stages of primary macrophage infection, we revealed critical insights into the dynamics of *S.* Typhimurium gene regulation. Our data revealed that the most abundant transcriptional changes occur at the onset of infection, with gene expression stabilizing as the infection progresses. This transition reflects the bacteria’s rapid adaptation to the evolving macrophage environment, characterized initially by immune-driven stress and later by metabolic exploitation of host-derived nutrients

### Metabolic adaptations underpinning intramacrophage survival

One of the key findings of our study is the identification of metabolic pathways supporting *S*. Typhimurium’s adaptation within macrophages. The upregulation of pathways for ethanolamine and propanediol metabolism underscores their importance as energy sources in the hypoxic macrophage niche. These findings expand on previous observations by demonstrating the critical role of host-derived nutrients in bacterial fitness under anaerobic conditions [37,38]. Moreover, our data highlight the activation of alternative electron acceptors, such as fumarate, tetrathionate, nitrate, and dimethyl sulfoxide, which collectively support energy production in oxygen-limited environments [68–71]. Interestingly, the observed downregulation of reactive oxygen species (ROS)-detoxifying enzymes after the early infection stage suggests a metabolic shift towards glycolysis and alternative antioxidant defenses, emphasizing the metabolic flexibility of *S.* Typhimurium in countering oxidative stress [72,73]. This confirms and extends previous findings regarding *S.* Typhimurium’s ability to exploit metabolic resources offered by macrophages at different times during infection [23,74].

### Novel role of the Psp system in intramacrophage survival

Our study provides the first direct evidence that the Psp system mediates *S.* Typhimurium resistance to antimicrobial peptides, specifically CRAMP, within macrophages. Previous work suggested that PspA helps maintain energy homeostasis in bacterial cells, thereby indirectly supporting the function of divalent metal transporters required to compete against host Nramp1 [52]. Nevertheless, the upregulation of the Psp system in host cells lacking Nramp1 [13,14] suggested an alternative function. We demonstrated that the Psp system is essential for resisting CRAMP and other inner membrane destabilizing agents. This represents a significant advancement in understanding the Psp system’s role in bacterial pathogenesis, addressing a longstanding question regarding its contribution to *S.* Typhimurium fitness in *Nramp1*-lacking hosts. Our results also reveal that the loss of PspA renders *S.* Typhimurium significantly more susceptible to CRAMP-induced killing. This susceptibility was prevented in macrophages deficient in CRAMP production (*Camp*^-/-^), conclusively linking PspA to resistance against this host antimicrobial peptide. The specificity of this function, combined with our evidence for the regulatory evolution of this system for intracellular expression by the SsrA-SsrB TCS, underscores the importance of the Psp system as a specialized AMP defense mechanism within the macrophage niche.

Future work is required to determine the precise mechanism by which PspA, and potentially other Psp proteins, contribute to antimicrobial peptide resistance in intracellular *S.* Typhimurium. In *E. coli,* current evidence suggests that to mitigate inner membrane damage and maintain the PMF, PspA assembles into large oligomers that directly bind the inner surface of the cytoplasmic membrane. The resulting scaffold reduces inner membrane permeability which may also block proton leakage [51,75,76]. Given that PspA, along with PspF, are the most conserved Psp proteins within *Enterobacteriaceae* and constitute the Psp minimal system [77], it is reasonable to postulate that PspA might function in a similar fashion in *S.* Typhimurium. Stabilization of the inner membrane by PspA oligomers would help mitigate damage caused by CRAMP. Alternatively, it is also possible that the reinforced cytoplasmic membrane could prevent the entry of CRAMP into the cytoplasm. Exploring these hypotheses should be the focus of future research as it will bring further insight into the role and importance of the Psp system in *S.* Typhimurium pathogenesis.

### Regulation of the Psp System by SsrA-SsrB

A surprising discovery of our study is the regulatory connection between the Psp system and the SsrA-SsrB two-component system (TCS) in *S.* Typhimurium. While PhoP-PhoQ and PmrA-PmrB are traditionally regarded as the primary regulators of antimicrobial peptide resistance in *S.* Typhimurium [43,78], we found that these TCSs do not regulate the *psp* operon. Instead, SsrA-SsrB, a master regulator of intracellular survival [79,80], directly controls *psp* expression in *S.* Typhimurium through an evolved regulatory sequence in the intergenic region upstream of *pspA* that differs in the non-pathogenic *S. bongori*. This novel regulatory relationship highlights the adaptability of the SsrB regulon, which has evolved to capture a large number of ancestral and horizontally-acquired genes critical for bacterial fitness in the intracellular environment [48,81,82].

In summary, our study advances the understanding of the temporal transcriptional landscape of *S.* Typhimurium during macrophage infection, identifying novel metabolic adaptations and unveiling the critical role of the Psp system in resistance to antimicrobial peptides. These findings provide a new perspective on how *S.* Typhimurium balances immune evasion and metabolic adaptation to thrive within host macrophages. Future research into the interplay between host defenses and bacterial stress responses will be instrumental in developing therapeutic strategies targeting either pathogen adaptations or macrophage immunity.

## Materials and methods

### Bacterial strains and growth conditions

A detailed list of the strains and plasmids used in this study is provided in Table 1. All experiments were performed with *S.* Typhimurium *enterica* serovar Typhimurium strain SL1344. Routine propagation of bacteria was in LB media (10 g/L NaCl, 10 g/L Tryptone, 5 g/L yeast extract) supplemented with appropriate antibiotics (streptomycin, 100 μg/mL; chloramphenicol, 34 μg/mL; ampicillin, 200 μg/mL). Where indicated, bacteria were grown in LPM media (5 mM KCl, 7.5 mM (NH4)2SO4, 0.5 mM K2SO4, 80 mM MES pH 5.8, 0.1% (w/v) casamino acids, 0.3% (v/v) glycerol, 24 μM MgCl2, 337 μM PO4^3-^). Bacteria were grown at 37°C with shaking.

**Table 1.** Bacterial strains used in this study.

| Strain | Source or reference |
| --- | --- |
| Wild-type *S. Typhimurium* strain SL1344 | [83] |
| Δ*pspA* *S. Typhimurium* strain SL1344 | This study |
| Δ*pspABC* *S. Typhimurium* strain SL1344 | This study |
| Δ*pspA* *S. Typhimurium* strain SL1344 + pGEN-MCS-*pspA* | This study |
| Wild-type *S. Typhimurium* strain SL1344 + pGEN-MCS | This study |
| Δ*ssrB* *S. Typhimurium* strain SL1344 | [47] |
| Δ*phoP* *S. Typhimurium* strain SL1344 | Coombes lab collection |
| Δ*pmrA* *S. Typhimurium* strain SL1344 | [84] |
| TOP10 *E. coli* | Invitrogen |

### Mice

Animal experiments were conducted according to Canadian Council on Animal Care guidelines using protocols approved by the Animal Review Ethics Board at McMaster University under Animal Use Protocol #20-12-41. Six to eight-week-old female C57BL/6J mice were purchased from Jackson Laboratories (000664) and C57BL/6 cathelicidin-null (*Camp*^-/-^) mice were provided by E. Cobo (University of Calgary). Animals were housed in a specific pathogen-free barrier unit under Level 2 conditions. Mice were fed a regular chow diet *at libitum*.

### Cell culture maintenance

Cells were maintained in a humidified incubator at 37°C with 5% CO2. Bone marrow-derived macrophages (BMDMs) were differentiated from the marrow isolated from the femur and tibia of female C57BL/6 or *Camp^-/-^* mice and maintained in RPMI containing 10% FBS (Gibco), 10% L929 fibroblast conditioned media, and 100 U penicillin-streptomycin. Cells were differentiated for 7 days in 150 mm Petri dishes, then lifted with ice-cold PBS for seeding in tissue culture-treated plates 20 h prior to infection. L929 fibroblast conditioned media was collected from the supernatants of L929 fibroblasts grown in DMEM with 10% FBS for 10 days.

### Cloning and mutant generation

Primers for cloning and mutant generation are listed in Table 2, and plasmids used in this study are listed in Table 3. All DNA manipulation procedures followed standard molecular biology protocols. Primers were synthesized by Sigma-Aldrich. PCRs were performed with Phusion, Phire II, or Taq DNA polymerases (ThermoFisher). All deletions and plasmid constructs were confirmed by PCR and verified by Sanger sequencing (McMaster Genomics Facility) or Whole Plasmid Sequencing (Plasmidsaurus, Oxford Nanopore Technology). In-frame, marked mutants of SL1344 *pspA* and *pspABC* were generated using Lambda Red Technology [85]. Wild-type SL1344 carrying pKD46 was transformed with linear PCR products containing gene-specific regions of homology and flanking the *cat* cassette carried by pKD3. Transformants were selected on LB agar supplemented with chloramphenicol (34 μg/mL) and knockouts were verified by PCR. To generate the *pspA* mutant complementation construct, the coding sequence of *pspA* and 800 bp upstream of the start codon was PCR-amplified and product was then cloned into pGEN-MCS after digesting with EcoRI/NotI (ThermoFisher). The sequenced-verified was transformed into Δ*pspA* S. Typhimurium strain SL1344 for expression. The transcriptional reporter constructs for P*pspABCDE* were generated by introducing the *pspABCDE* promoter into a luciferase reporter plasmid as previously described [86]. Briefly, ∼500 bp regulatory region upstream of *pspABCDE* from SL1344 was PCR-amplified and cloned into the BamHI/SnaBI-digested pGEN-*luxCDABE* plasmid. The sequenced-verified plasmid was transformed into electrocompetent wild-type SL1344 or derivate strains.

**Table 2.** Plasmids used in this study.

| Plasmid | Description | Reference |
| --- | --- | --- |
| pKD3 | Template plasmid for Lambda Red recombination | [85] |
| pKD46 | Lambda Red recombinase expression plasmid | [85] |
| pGEN-*luxCDABE* | Lux transcriptional reporter plasmid | [87] |
| pGEN-MCS | Low-copy-no. cloning vector | [87] |
| pGEN-P*pspABCDE-luxCDABE* | Lux transcriptional reporter for *pspABCDE* promoter | This study |
| pGEN-P*pspABCDE-S.bongori-ssrB-luxCDABE* | Lux transcriptional reporter for *pspABCDE* promoter with putative SsrB-binding site from *S. bongori* | This study |
| pGEN-P*pspABCDE-scrambled-ssrB-luxCDABE* | Lux transcriptional reporter for *pspABCDE* promoter with scrambled putative SsrB-binding site | This study |
| pGEN-MCS-*pspA* | *pspA* with 800 bp upstream of coding sequence cloned into pGEN-MCS for complementation experiments | This study |

**Table 3.**
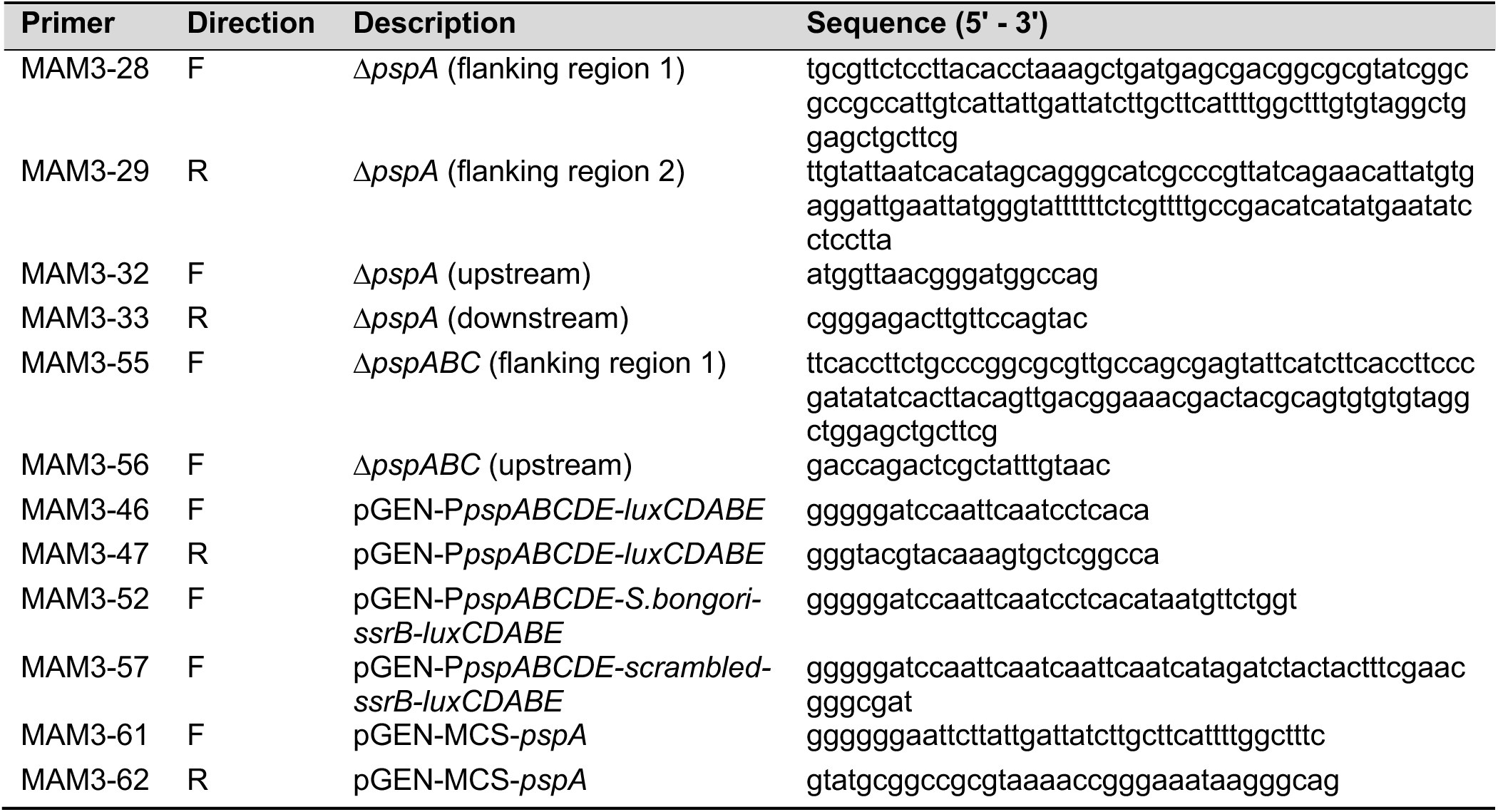
Primers used in this study.

### RNA isolation of intracellular bacteria for sequencing

Wild-type SL1344 was enriched from infected bone marrow-derived macrophages (C57BL/6J) as previously described with modifications [16]. Macrophages were seeded 20 h prior to infection at ∼10^7^ cells/100 cm^2^ tissue culture dish in RPMI containing 10% FBS (Gibco) with 100 ng/mL LPS from *S. enterica* serovar Minnesota R595 (Millipore). Overnight cultures of wild-type SL1344 were diluted to obtain a multiplicity of infection ∼50:1 and added to each plate. Plates were spun down at 500 x *g* for 2 min, then incubated for 20 min at 37°C with 5% CO2. Media was aspirated, plates were washed 3 times with PBS, and fresh RPMI with 10% FBS and 100 μg/mL gentamicin to kill extracellular bacteria was added. After a second incubation of 30 min at 37°C with 5% CO2, media was aspirated and replaced with fresh RPMI with 10% FBS and 10 μg/mL gentamicin. At this time (0 h) or 4 h, 8 h, and 12 h later, media was aspirated, plates were washed twice with PBS, then infected macrophages were lysed on ice for 20 min in ice-cold 0.2% SDS, 1% acidic phenol, 19% ethanol in DEPC water. Lysates containing intracellular bacteria were collected and centrifuged at 4,000 rpm for 10 min, 4°C. After three washes in ice-cold wash buffer (1% acidic phenol, 19% ethanol in DEPC water) (4,000 rpm, 10 min, 4°C each wash), the supernatant was discarded, the bacterial pellet was resuspended in the remaining liquid, transferred to a clean RNAse-free microcentrifuge tube, then stored at - 80°C. RNA was extracted using a MasterPure RNA purification kit (Epicentre) and treated with DNase I (Invitrogen). The quality of the purified RNA was verified with an Agilent RNA 6000 Pico BioAnalyzer.

### Sequencing, mapping of RNA-seq libraries, and differential gene expression analysis

All RNA-seq data are from three independent experiments per time-point. Prior to RNA-sequencing, ribosomal RNA was depleted using Ribo-Zero (Illumina), followed by the generation of barcoded cDNA for each sample. cDNA was sequenced on an Illumina NextSeq P2 platform with single-end reads. Raw reads were processed using FastQC [88] and trimmed with Trimmomatic [89] to remove Truseq adapter sequences. Sequencing data was mapped to the host reference genome GRCm39 (GCF_000001635.27) using BWA (mem algorithm) with default settings to remove non-bacterial sequences, then mapped to the reference genome of *S.* Typhimurium SL1344 (NC_016810) using BWA with default settings [90]. Uniquely-mapped reads were quantified using FeatureCounts [91] and the differential gene expression was determined using the R package DESeq2 [26]. The Likelihood ratio test (LTR) and reduced model parameters were used to identify genes that show changes in expression across all time points. Genes were deemed differentially regulated if they showed an FDR-adjusted *P* value <

0.1. For clustering analysis, regularized log transformation of the normalized counts of the differentially regulated genes were clustered using degPatterns from the R package DEGreport. The function was run using the default parameters and the produced clusters were visualized using the R package degPlotCluster [92]. Clusters of Orthologous Gene (COG) were assigned to significantly regulated genes using COGclassifier 1.0.5 Python package [93]. DESeq2 using the default parameters was used to conduct differential expression analysis of *S.* Typhimurium genes at early, middle, and late stages of infection relative to the onset stage. Genes were considered differentially regulated if they showed log2 fold change > 1 or < −1 at FDR-adjusted P value < 0.01. Circular visualization of the data was generated using the R package Circlize [94]. To generate the heatmap, transcripts per million (TPM) values of the significantly regulated genes were calculated for each time point and values were then used to generate the heatmap using Pheatmap R package [95]. RNA-sequencing data that support the findings of this study have been deposited in the National Center for Biotechnology Information Gene Expression Omnibus (GEO) and are accessible through the GEO Series accession number GSE294365.

### Transcriptional reporter assays

Strains containing pGEN-*lux* promoter-reporter plasmid were grown in LB until the mid-log phase, then subcultured 1:50 into LPM media in black 96-well flat, clear-bottom plates (Corning). Plates were incubated at 37°C with shaking, and luminescence and OD600 were measured at 30-minute intervals for 16 h using an Agilent BioTek Cytation 5. Luminescence (RLU) was normalized to OD600.

### Intracellular replication assays

Differentiated bone marrow-derived macrophages (C57BL/6 or *Camp*^-/-^) were seeded 20 h prior to infection at 10^5^ cells/well in 96-well tissue culture plates in RPMI containing 10% FBS (Gibco) with 100 ng/mL LPS from *S. enterica* serovar Minnesota R595 (Millipore). Overnight cultures of wild-type SL1344 or Δ*pspA* mutant were diluted to obtain a multiplicity of infection ∼10:1 and added to each well. Infected plates were spun for 2 min at 500 x *g* and incubated at 37°C with 5% CO2. Following 30 min of infection, bacteria-containing media was removed, macrophages were washed 3 times with PBS, and fresh RPMI with 10% FBS and 100 μg/mL gentamicin was added to each well to eliminate extracellular bacteria. Plates were incubated again for 30 min at 37°C with 5% CO2. Gentamicin-containing media was removed from macrophages and replaced with fresh RPMI with 10% FBS and 10 μg/mL gentamicin. Immediately after this media replacement step, adhered macrophages from 1/5 of the wells were washed twice with PBS and lysed in sterile water. Bacterial colony-forming units (CFUs) from each lysed well were enumerated by serially diluting in PBS and plating on LB plates supplemented with 34 μg/mL of chloramphenicol for Δ*pspA* (CFU at 0 h). After 4 h, 8 h, 12 h, or 24 h of incubation at 37°C with 5% CO2, adhered macrophages from remaining wells were washed with PBS and lysed in sterile water for plating and CFU enumeration. Fold change in CFU (CFU at 4 h, 8 h, 12 h, or 24 h divided by 0 h) was calculated to represent replication throughout the experiment.

### Competitive infection

For bacterial preparation, wild-type SL1344 or Δ*pspABC* mutant strains were grown overnight in LB medium with appropriate antibiotic selection. For competitive infections, the inoculum consisted of a 1:1 ratio of wild-type strain resistant to streptomycin and a second competing mutant strain additionally resistant to chloramphenicol. Mice were infected intraperitoneally (IP) with 2 x 10^5^ CFU bacteria in 0.1 M HEPES (pH 8.0) with 0.9% NaCl. After 45 hours mice were sacrificed, and total bacterial loads in the cecum, spleen, and liver were enumerated from organ homogenates serially diluted and plated on LB medium containing streptomycin. Colonies were replica-plated with chloramphenicol selection for the enumeration of mutant CFU. The competitive index was calculated using the following formula:

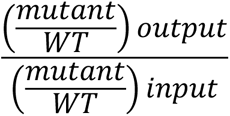

### Minimum inhibitory concentration assays

MIC determinations for PMB, colistin, CCCP, EDTA, polymyxin B nonapeptide, and CRAMP were performed using broth microdilution in 96-well flat, clear-bottom plates (Corning). Compounds were serially diluted two-fold starting at the following concentration: PMB and colistin; 100 μM, CCCP; 80μM, EDTA; 2 μM, PMBN, 50 μM, and CRAMP; 33 μM. Bacterial cultures grown overnight in LB were diluted ∼1: 5 000 into LPM, then added to the compound-containing media. OD600 was read immediately after bacteria addition (OD0h) and after ∼20 h of incubation at 37°C (OD20h) using a PerkinElmer plate reader. For the CRAMP MIC curve, percentage growth was calculated by subtracting OD0h from OD20h, then normalizing values to a water control set to 100% growth.

### CRAMP, hydrogen peroxide, and spermine NONOate survival assays

Strains grown overnight in LB were subculture 1:50 into LPM and incubated at 37°C with shaking for 3 h. Bacteria were harvested and normalized to an OD600 of 0.5. Cells were washed and resuspended in LPM and diluted 1:10 (OD600 of 0.005). Each strain was incubated in LPM containing 10 μM of CRAMP, 50 μM of H_2_O_2_, or 50 μM of Spermine NONOate for 3 h at 37°C. The number of viable bacteria was determined by plating on LB agar supplemented with 34 μg/mL of chloramphenicol for Δ*pspA* and *ΔphoP* strains, and percentage survival was calculated as the number of CFU/mL at 3 h relative to 0 h.

### Data and statistical analysis

Data were analyzed using RStudio version 2024.04.2+764 with R version 4.4.1, and GraphPad Prism 10.1.1 software (GraphPad Inc., San Diego, CA). A *P* value of 0.05 or less was considered significant. An explanation of the software used for RNA-sequencing analysis can be found in the corresponding experimental method description.

## Acknowledgments

We thank the McMaster Genome Facility for performing RNA-sequencing, and E. Cobo for providing the *Camp*^-/-^ mice. We are grateful to M. Zangara for assistance with BMDMs isolation, and to members of the Coombes lab for helpful discussions on this work.

## Supporting information

**S1 Table.** Summary of the data from the clustering analysis of the RNA-sequencing. DESeq2 performs a LTR test and reports P values that are adjusted for multiple testing using the procedure of Benjamini and Hochberg. Genes showing values showing adjusted *P* value < 0.01 were considered significant. Clustering was performed using degPatterns from the R package DEGreport. COG assignment data of all genes are reported as identified by COGclassifier. Cells with NA indicate that the gene could not be assigned to a particular COG family. RNA-sequencing data that support the findings of this study have been deposited in the National Center for Biotechnology Information Gene Expression Omnibus (GEO) and are accessible through the GEO Series accession number GSE294365

**S2 Table.** Summary of the data from the differential gene expression analysis of the RNA-sequencing. DESeq2 performs a Wald test and reports *P* values that are adjusted for multiple testing using the procedure of Benjamin and Hochberg. Genes showing values > 2 or < -2 for log2 fold change in expression at early, middle and late stages of infection relative to onset at adjusted *P* value < 0.01 were considered significant.

**S1 Fig.**
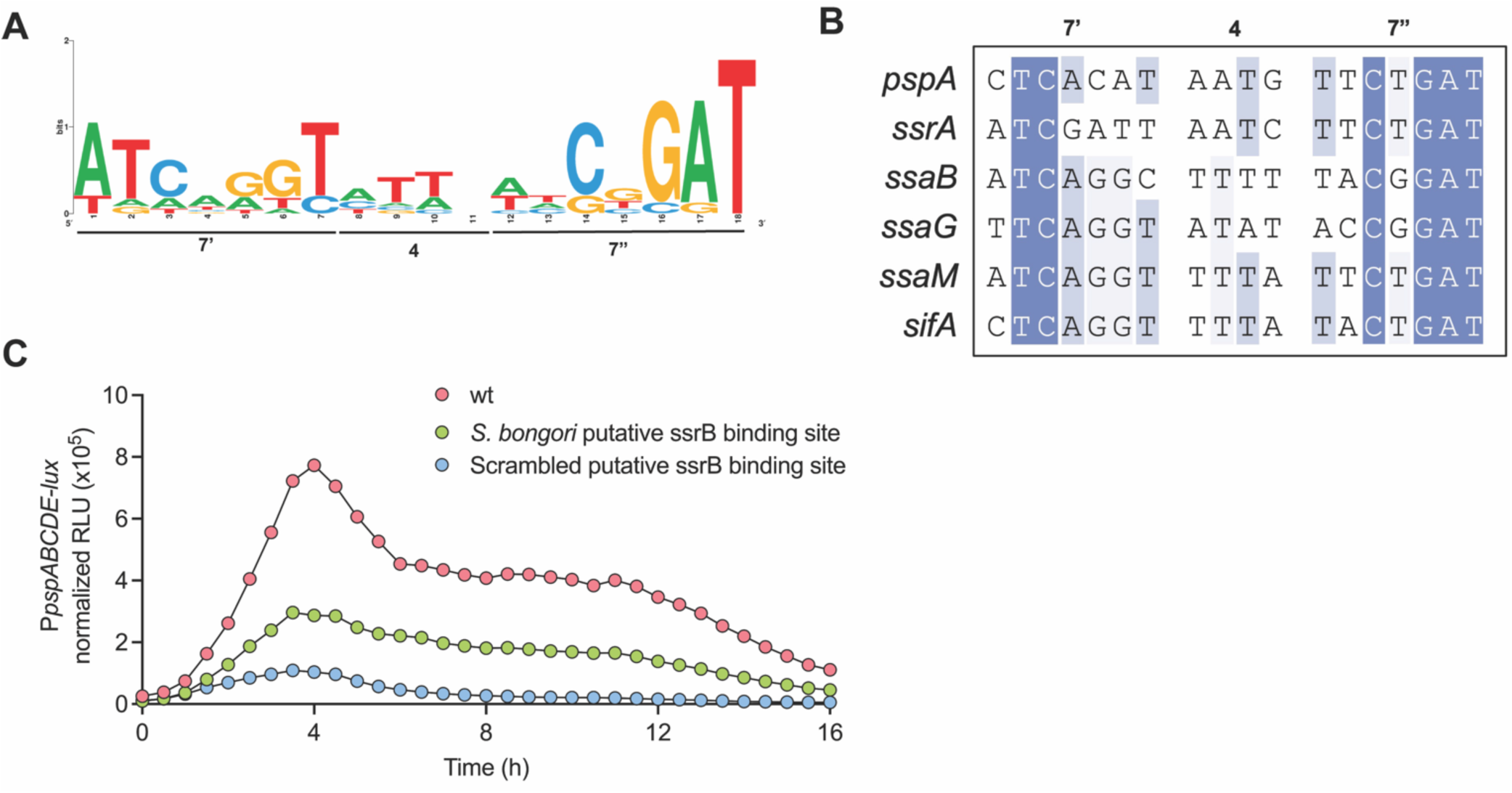
Identification of the SsrB putative binding site upstream of the *psp* operon. (A) Consensus motif logo of SsrB-binding site identified by Tomljenovic-Berube, *et. al.* (2013). (B) Aligned putative SsrB-binding site upstream of the *pspA* gene and known SsrB-regulated SPI-2 genes in *S.* Typhimurium. The left (7′) and right (7″) heptamers and the 4-bp spacer are displayed as a heat map to show bases of high conservation (dark blue) from degenerate regions (light blue/white). (C) Transcriptional reporter assay of the wild-type (wt) P*pspABCDE*- *lux* and the two mutated SsrB-binding site (putative *S. bongori* and scrambled SsrB-binding site) in wild-type *S.* Typhimurium grown in LPM for 16h. Data are mean relative light units (RLU) normalized to the optical density of the culture from three independent experiments.

**S2 Fig.**
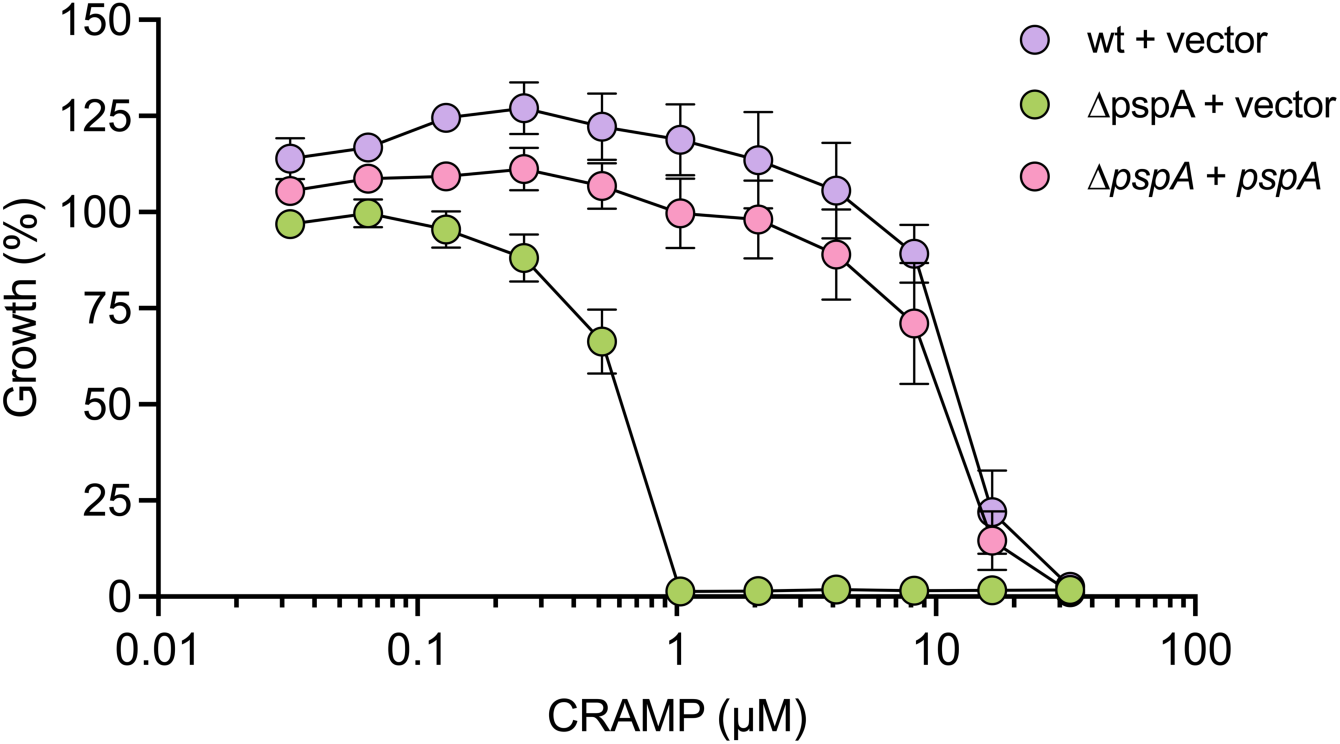
Complementation with *pspA* restores resistance to CRAMP. Growth in the presence of CRAMP in infection-mimicking media of *S.* Typhimurium Δ*pspA* mutant complemented with *pspA*, and strains carrying the empty pGEN-MCS vector control. Data are from at least two biological replicates, dots and error indicate mean and standard error of the mean.

